## Supplementary Table and Figure for "Micro-dissection and integration of long and short reads to create a robust catalog of kidney compartment-specific isoforms"

Table S1. Compartment-specific genes expression enrichment analysis.

| **Glomerular-only expressed genes** | |
|---|---|
| **KEGG Pathway** | **P-Value** |
| Rap1 signaling pathway | 2.90E-06 |
| Ribosome | 8.30E-06 |
| Non-alcoholic fatty liver disease (NAFLD) | 1.10E-04 |
| Thyroid hormone signaling pathway | 1.40E-04 |
| Transcriptional misregulation in cancer | 1.90E-04 |
| Neurotrophin signaling pathway | 2.50E-04 |
| Epstein-Barr virus infection | 3.70E-04 |
| TNF signaling pathway | 5.10E-04 |
| Wnt signaling pathway | 5.50E-04 |
| Pathways in cancer | 6.80E-04 |
| Adherens junction | 8.80E-04 |
| Platelet activation | 9.10E-04 |
| Bacterial invasion of epithelial cells | 9.50E-04 |
| Colorectal cancer | 1.10E-03 |
| Ubiquitin mediated proteolysis | 1.70E-03 |
| Osteoclast differentiation | 2.00E-03 |
| Alzheimer's disease | 2.10E-03 |
| Herpes simplex infection | 2.10E-03 |
| Proteoglycans in cancer | 2.70E-03 |
| Endocytosis | 2.90E-03 |
| HIF-1 signaling pathway | 3.50E-03 |
| Parkinson's disease | 3.70E-03 |
| Sphingolipid signaling pathway | 3.70E-03 |
| Spliceosome | 5.10E-03 |
| mTOR signaling pathway | 5.30E-03 |
| Vascular smooth muscle contraction | 5.70E-03 |
| Viral carcinogenesis | 8.00E-03 |
| Chemokine signaling pathway | 8.30E-03 |
| Leukocyte transendothelial migration | 8.70E-03 |
| Oxidative phosphorylation | 8.70E-03 |
| Viral myocarditis | 9.30E-03 |
| MAPK signaling pathway | 1.10E-02 |

| **Tubular-interstitial-only expressed genes** | |
|---|---|
| **KEGG Pathway** | **P-Value** |
| Metabolic pathways | 4.80E-07 |
| Ubiquitin mediated proteolysis | 9.90E-05 |
| Aldosterone-regulated sodium reabsorption | 2.70E-04 |

| | |
|---|---|
| GnRH signaling pathway | 7.60E-04 |
| Gastric acid secretion | 7.90E-04 |
| Axon guidance | 1.10E-03 |
| Sphingolipid signaling pathway | 1.20E-03 |
| Neurotrophin signaling pathway | 1.20E-03 |
| Renal cell carcinoma | 1.60E-03 |
| Oxytocin signaling pathway | 1.90E-03 |
| Insulin resistance | 2.30E-03 |
| Choline metabolism in cancer | 2.60E-03 |
| Central carbon metabolism in cancer | 2.90E-03 |
| Inflammatory mediator regulation of TRP channels | 3.00E-03 |
| Aldosterone synthesis and secretion | 4.50E-03 |

Figure S1. Distribution of the lengths of the consensus full-length transcripts.

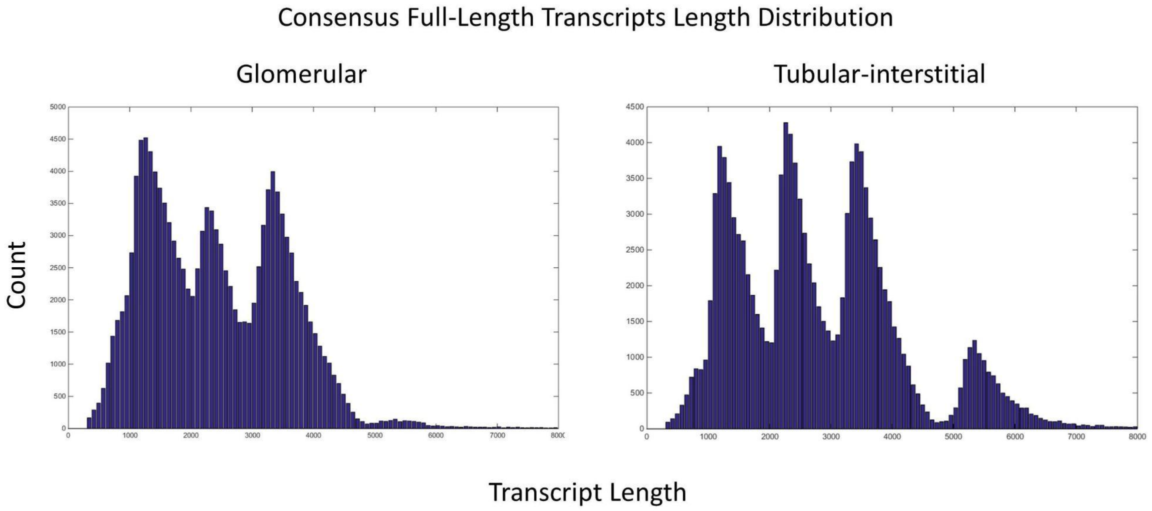
